# Nose-to-Brain Administration of Cannabidiol-Loaded Polymeric Micelles Improves the Core Behavioral Symptoms of Autism Spectrum Disorder

**DOI:** 10.1101/2025.11.05.686695

**Authors:** Rania Awad, Shlomit Aga-Mizrachi, Inon Maoz, Lilach Simchi, Avi Avital, Alejandro Sosnik

## Abstract

Neurodevelopmental disorders including autism spectrum disorder (ASD) affect 5.9% of the global population. Research shows the potential therapeutic use of cannabidiol (CBD) to treat different neurodevelopmental disorders, including ASD. Intranasal drug delivery (i.n.) is a non-invasive and painless administration route that enhances drug bioavailability in the brain by bypassing the blood-brain barrier. However, i.n. has limited bioavailability due to the low nasal mucosa permeability. Various polymeric nanoparticles have been investigated for i.n. delivery with different successes. In this study, we developed and characterized polymeric micelles of the poly(ethylene oxide)-*b*-poly(propylene oxide) block copolymer Pluronic^®^ F127 loaded with 25% w/w CBD for nose-to-brain delivery in ASD. CBD-loaded polymeric micelles display a hydrodynamic diameter of 41 ± 1 nm by Intensity and 23 ± 1 nm by Number, as measured by dynamic light scattering, and showed very good compatibility and permeability in the human nasal septum cell line RPMI 2650, an *in vitro* model of the nasal epithelium. The accumulation of CBD-loaded polymeric micelles upon i.n. administration to autistic rats is confirmed by bioimaging. The pharmacokinetics of CBD upon i.n. (dose of 5 mg/kg) and oral (15 mg/kg) administration of the loaded polymeric micelles shows a 27.8% increase of the CBD concentration in the brain 20 min of autistic rats after i.n. administration, despite the 3-fold decrease in the dose. Finally, the efficacy of this nanoformulation to improve the core symptoms of ASD is demonstrated in behavioral studies in a rat model of the disorder.

## 1. Introduction

Autism Spectrum Disorder (ASD) is a heritable, heterogeneous neurodevelopmental disorder characterized by impaired social interaction and communication, repetitive behaviors, deficit in neurocognitive development, anxiety, emotional dysregulation and disrupted sleeping and eating habits.^1,2,3^ According to the World Health Organization, the global prevalence of ASD is one in 100 children,^4,5^ and in the US is one in 36 children.^6^ The ASD diagnosis is challenging owing to its complex clinical manifestation, requiring a careful behavioral assessment. ASD is associated with alterations in some regions of the central nervous system (CNS), especially the limbic system, including the hippocampus, amygdala, and cerebellum.^7^ Currently, there is no clinically proven pharmacological treatment to tackle the core symptoms of ASD,^8^ and different approved psychiatric drugs have been repurposed to decrease some of the core symptoms such as seizures, anxiety, depression, extreme emotional reactions and obsessive behavior.^9,10,11^ In this scenario, there is an urgent need to investigate new pharmacological treatments to improve the quality of life of autistic people.

Cannabidiol (CBD) is a non-psychoactive cannabinoid isolated from the *Cannabis sativa* plant that has been proposed as a potential treatment for various conditions, including pain, inflammation and CNS conditions such as ASD.^12,13,14^ CBD interacts with the endocannabinoid system (ECS), which is a lipid-based signaling network involved in CNS development and immune regulation, that has been increasingly implicated in the pathophysiology of ASD.^12,15^ Cannabinoid receptors CB1 and CB2 are differentially expressed in key brain regions affected in ASD and in immune cells, respectively. Dysregulation of ECS components, such as altered CB2 receptor expression and elevated microglial activation, has been observed in both ASD patients and animal models, suggesting a role in neuroinflammation and behavioral disturbances. CBD has emerged as a promising therapeutic agent due to its ability to modulate ECS activity, particularly by increasing anandamide levels through inhibition of fatty acid amide hydrolase 1, the enzyme responsible for its metabolism.^12,16,17,18^ Several studies showed that CBD can reduce autistic-like behaviors, and improve social interactions.^19,20^ In ASD, CBD is often administered as CBD-enriched cannabis extracts.^21,22^ However, CBD exhibits high lipophilicity (log*P* = 6.3) and poor aqueous solubility (∼10 mg/L),^23^ which result in very low oral bioavailability that limits its therapeutic efficacy;^24^ the oral bioavailability of CBD in blood and brain is 6-15% and 1-3%, respectively.^12^ The nanoencapsulation of CBD within polymeric and lipidic nanocarriers has been proposed to increase its permeability across biological barriers by different administration routes.^12,15,25,26,27^

Intranasal (i.n.) drug delivery is a noninvasive, painless, self-administration route that enables the fast and direct transport of nanoparticles deposited in the olfactory epithelium of the nasal mucosa to the brain, bypassing the blood-brain-barrier (BBB) and hepatic first-pass metabolism, which enhances drug bioavailability and minimizes systemic side effects.^28,29^ A plethora of cellular mechanisms are implicated in the nose-to-brain transport of nanoparticulate matter.^28,30,31^ Intranasally administered nanoparticles face a series of mucosal barriers before their transport to the brain takes place, including an outer mucus layer, nasal epithelial tight junctions and mucociliary clearance, and a mucoadhesive and muco-penetrating nanoparticles have been developed to take advantage of this delivery pathway.^28^ For example, nanoencapsulation of the antiretroviral efavirenz within polymeric micelles of linear and branched poly(ethylene oxide)-*b*-poly(propylene oxide) (PEO-PPO) block copolymers^32^ showed a statistically significant increase in the brain bioavailability to intravenous injection.^33^

Aiming to lead to a breakthrough in the therapeutic use of CBD for ASD, in this work, we design and comprehensively characterize CBD-loaded polymeric micelles with a high payload of up to 27% w/w. After demonstrating the ability of the nanoparticles to cross a model of the nasal epithelium *in vitro*, we comparatively investigate the CBD pharmacokinetics in blood and brain after i.n. and oral administration. Finally, pharmacodynamically, we demonstrate the improvement of social cooperation (SC) and other core ASD symptoms in a behavioral rat model.^34^

## 2. Experimental section

### 2.1. Preparation and characterization of CBD-loaded Pluronic^®^ F127 polymeric micelles

To prepare CBD-loaded 10% w/w polymeric micelles, Pluronic^®^ F127 (1 g, kindly donated by BASF, Vandalia, IL, USA) was dissolved in Milli-Q water (9 g, Mili-Q^®^ Direct 8, Millipore SAS, Molsheim, France) and left at room temperature (RT) for at least 24 h. To assess the encapsulation capacity, various amounts of CBD (0.049-0.291 g, CBDepot.eu, Teplice, Czech Republic) were added to the 10% w/w polymeric micelles (9.737-9.7 g) and the mixture magnetically stirred (300 RPM) at RT for 48-72 h protected from light to prevent the photodegradation of cargo. Then, the system was filtered (0.2 μm Minisart^®^, Sartorius Stedim Biotech SA, Göttingen, Germany) to remove any insoluble CBD, and the CBD concentration was quantified by the Beam’s test. Briefly, a solution of 5% w/v KOH (Bio-Lab Ltd., Jerusalem, Israel) in absolute ethanol (Gadot, Netanya, Israel) was prepared and kept at RT until use. Then, the corresponding sample for analysis was dissolved in absolute ethanol, 5% w/v KOH solution (∼250 µL) was added, and the sample was stirred for 5 min to allow the oxidation of the extracted CBD to HU-331 and bis-HU-331. Then, the absorbance was measured at 330 nm in a Multiskan GO Microplate Spectrophotometer (Thermo Fisher Scientific Oy, Vantaa, Finland) and interpolated in a calibration curve of CBD in absolute ethanol with concentrations between 0.0001% and 0.1% w/v (R^2^ = 0.9992), treated with 5% w/v KOH and the CBD concentration calculated.^22^ For all the subsequent experiments, a CBD loading of 25% w/w (0.243% w/v final CBD concentration in the nanosuspension) was selected to ensure complete encapsulation and long-term physical stability. Nanosuspensions were stored at 4°C until use. For some analyses, samples were frozen in liquid nitrogen and freeze-dried using a Labconco Free Zone 4.5 plus L Benchtop Freeze Dry System (Labconco, Kansas City, MO, USA).

The size (expressed as hydrodynamic diameter, D_h_) and polydispersity index (PDI, a measure of the size distribution) were measured using a Malvern Zetasizer-Nano (ZEN 3600, Malvern Instruments, Malvern, UK) equipped with a 633 nm laser at a scattering angle of 173°, at 25°C. Data was analyzed using CONTIN algorithms (Malvern Instruments). For zeta-potential (Z-potential) measurements, the laser Doppler micro-electrophoresis setup in the same instrument was utilized. Values are expressed as mean ± S.D. of three independent samples prepared under the same conditions.

The thermal behavior of CBD in the polymeric micelles was compared to pure CBD and pristine Pluronic^®^ F127 by differential scanning calorimetry (DSC) using a Mettler Toledo DSC-1 STAR system (Metter-Toledo, Schwerzenbach, Switzerland). For this, freeze-dried samples (5-10 mg) were sealed in 40 µL Al-crucibles pans (Metter-Toledo) and heated from −70 to 250°C (10°C/min). To assess the amorphous or crystalline nature CBD in the polymeric micelles, freeze-dried samples were analyzed by power X-ray diffraction (PXRD, SmartLab 9 kW, Rigaku Corp., Tokyo, Japan). Measurements were conducted in a 2θ range of 5–60◦ using Cu-Ka radiation (λ = 0.15406 nm).

The pH of 25% w/w CBD-loaded Pluronic^®^ F127 polymeric micelles was measured with a Sartorius pH meter (Gottingen, Germany). The viscosity of the 25% w/w CBD-loaded Pluronic^®^ F127 polymeric micelles was measured using an Ostwald viscometer. Before using it, the viscometer was washed overnight in a 0.1 N solution of HCl (Sigma-Aldrich, St. Louis, MO, USA). The viscometer was then rinsed thoroughly with Milli-Q water and allowed to dry overnight in an oven, at 80°C. Once cooled to RT, the elution time of the liquids in the viscometer is measured, and the viscosity η1 is calculated with Equation 1

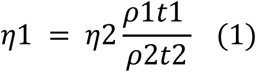

where ρ1 is the density of the nanoformulation, t1 is the time the nanoformulation takes to flow, under the influence of gravity, between two points scribed on the viscometer, η2 is the viscosity of Milli-Q water (1 mPa.s), ρ2 is the density of milli-Q water and t2 is the time it takes Milli-Q water to flow between the same two lines at the same temperature.

The morphology and nanostructure of 25% w/w CBD-loaded Pluronic^®^ F127 polymeric micelles were visualized by cryogenic transmission electron microscopy (cryo-TEM). Samples were vitrified in a controlled environment vitrification system (CEVS). For this, ∼3 µL of the sample was placed on a carbon-coated perforated polymeric film placed on a TEM grid (200 mesh), mounted on tweezers, and the sample turned into a thin film (thickness <300 nm) by blotting the excess of the solution with a filter paper-covered metal strip. The grid was plunged into liquid ethane and transferred and kept in liquid nitrogen. Samples were visualized in a FEI Talos 200C High Resolution TEM (Thermo Fisher Scientific, Waltham, MA, USA) equipped with a cryo-holder (cryo-specimen holder, Gatan, Inc., Pleasanton, CA, USA) at −181 °C and an accelerating voltage of 120 kV.

The visualization of the 25% w/w CBD-loaded polymeric micelles in aqueous dispersion and the subsequent quantification of their concentration (expressed in particles per mL) was carried out by nanoparticle tracking analysis (NTA) in a Malvern NanoSight^®^ NS500-Zeta HSB system with high sensitivity camera and red laser of 638 nm for fluorescent analysis (Malvern Instruments), at 37°C. The initial nanoparticle dispersion was diluted (1:5) in Milli-Q water before the analysis fit the measurement range of the NTA instrument and the particle concentration was corrected by the dilution factor.

### 2.2. CBD release in vitro

The release profile of CBD from the Pluronic^®^ F127 polymeric micelles was conducted using the dialysis membrane method. Briefly, a 25% w/w CBD-loaded nanosuspension (5 mL) was poured into dialysis membranes (regenerated cellulose dialysis membrane with molecular weight cut-off of 3500 Da, Cellu-Sup T1 nominal flat width of 46 mm, diameter of 29.3 mm, and volume/length ratio of 6.74 mL/cm, Membrane Filtration Products, Inc., Seguin, TX, USA) prewashed in Milli-Q water for 30 min. Then, samples were immersed in glass vials filled with simulated nasal electrolyte solution (SNES, 20 mL) comprising an aqueous solution containing 8.77 mg/mL sodium chloride (Sigma-Aldrich), 2.98 mg/mL potassium chloride (Sigma-Aldrich) and 0.59 mg/mL calcium chloride dihydrate (Sigma-Aldrich) of pH 7.4,^35^ sealed with parafilm and stirred under magnetic stirring (300 RPM) at 37°C. At predetermined time intervals, aliquots (500 µL) were withdrawn and replaced with an equal volume of fresh release medium. Then, the sample was treated with a 5% w/v KOH water solution (250 µL, Bio-Lab Ltd., Jerusalem, Israel), the absorbance was measured at 330 nm in a Multiskan GO Microplate Spectrophotometer (Thermo Fisher Scientific Oy, Vantaa, Finland), and interpolated in a CBD calibration curve. To build the calibration curve, a CBD stock was prepared by dissolving it in methanol (5% w/v, Gadot, Netanya, Israel), 1 mL of this stock solution was diluted in different volumes (3-10 mL) of SNES to obtain final CBD concentrations between 0.0001% and 0.25% w/v, and the solutions reacted with a 5% w/v KOH water solution (R^2^ = 0.998). Then, the cumulative CBD release (expressed in percentage of the initial amount) was calculated and plotted against time. Results are expressed as mean ± S.D. of at least three independent experiments. The release data were fitted to different models using the DDSolver software.^36^

### 2.3. Cell compatibility and permeability across model of the nasal epithelium in vitro

The compatibility of 25% w/w CBD-loaded Pluronic^®^ F127 polymeric micelles was assessed in the human nasal septum epithelium cell line RPMI 2650 (ATCC CMCL-30, American Type Culture Collection, Manassas, VA, USA).^37^ For this, the cells were cultured in 96-well plates (5 x 10^3^ cells/well) with Minimum Essential Medium (MEM) Eagle (Sigma-Aldrich) supplemented with L-glutamine, 10% heat-inactivated fetal bovine serum (Sigma-Aldrich), and a penicillin/streptomycin antibiotic mixture (5 mL of a commercial mixture of 100 U per mL penicillin + 100 μg per mL streptomycin per 500 mL medium, Sigma-Aldrich) and incubated at 37°C in a humidified 5% CO_2_ environment and split every 4−5 days. Then, the cell culture medium was replaced by 200 μL of medium containing 25% w/w CBD-loaded Pluronic^®^ F127 polymeric micelles (different CBD concentrations in range of 1.178-8.588 µM) and the cells incubated for 24 h. After 24 h, the medium was removed, and fresh pre-heated medium (100 μL) and sterile 3-(4,5-dimethylthiazol-2-yl)-2,5-diphenyltetrazolium bromide (MTT) solution (25 µL, 5 mg/mL, Sigma-Aldrich) was added. Cells were incubated for 3 h at 37°C under 5% CO_2_ environment, the supernatant removed, the formazan crystals dissolved in dimethyl sulfoxide (100 μL, DMSO, CARLO ERBA Reagents, Emmendingen, Germany) and the absorbance measured at 530 nm with reference at 670 nm (Multiskan GO Microplate Spectrophotometer). The percentage of live cells was calculated with respect to a control of cells incubated only with culture medium that were considered 100% viable. Results are expressed as mean ± S.D.

The apparent permeability of 25% w/w CBD-loaded Pluronic^®^ F127 polymeric micelles was evaluated in RPMI 2650 cell monolayers.^37^ For this, cells were cultured onto cell culture inserts (ThinCert, culture surface of 113.1 mm^2^, 3.0 μm pore size, Greiner Bio-One GmbH, Frickenhausen, Germany) in 12-well plates (15.85 mm diameter, 16.25 mm height, Greiner CELLSTAR, Monroe, NC, USA) with 0.5 and 1.5 mL of MEM in the apical and basolateral compartments, respectively. Cultures were maintained in a liquid-covered culture for eight days, and the medium was replaced every 2−3 day. After 8 days, inserts were exposed to air to generate an air-liquid interface (ALI) for 1−2 more weeks. The integrity of the cell monolayer was characterized by transepithelial electrical resistance (TEER) measurements performed with an epithelial volt-ohmmeter (“EVOM2”, WPI, Sarasota, FL, USA). For permeability experiments, only inserts that have resistance >150 Ω/cm^2^ were used and polymeric micelles were prepared with 80% w/w of pristine copolymer and 20% w/w of copolymer fluorescently labeled by the conjugation of fluorescein isothiocyanate (FITC, Sigma-Aldrich) to the terminal hydroxyl groups. Briefly, Pluronic^®^ F127 (100 mg) was dissolved in dry dimethyl formamide (2 mL, DMF, Bio-Lab Ltd.) under magnetic stirring; DMF was dried with activated molecular sieves 3A (Sigma-Aldrich) before use. FITC was dissolved in dry DMF (70 mg/mL) and the solution (0.2 mL) was added to the Pluronic^®^ F127 solution, under magnetic stirring and allowed to react for 16 h protected from light, at 32°C. Finally, the product was dialyzed (48 h, regenerated cellulose dialysis membranes, molecular weight cut-off of 3500 Da), frozen in liquid nitrogen, and freeze-dried (72−96 h, Labconco Free Zone 4.5 plus L Benchtop Freeze Dry System). The preparation of CBD-loaded Pluronic^®^ F127 polymeric micelles was conducted as described above though replacing 20% of the pristine copolymer in the formulation with FITC-labeled one. Then, FITC-labeled 25% w/w CBD-loaded Pluronic^®^ F127 polymeric micelles were diluted to a final copolymer concentration of 0.01% and 0.05 w/v in Hank’s Balanced Salt Solution (HBSS, Sigma-Aldrich) buffered to pH 7.2 with 25 mM 4-(2-hydroxyethyl)-1-piperazineethanesulfonic acid (HEPES, Sigma-Aldrich) that was used as transport medium. At the beginning of the experiment, the medium in the apical and basolateral compartments was replaced with transport medium (HBSS) and the cells were incubated for 15 min at 37°C in a humidified 5% CO_2_ atmosphere to enable their adjustment to the new medium. Then, the transport medium in the donor (apical) compartment was replaced by the corresponding sample (0.4 mL) and in the acceptor compartment (basolateral) by fresh transport medium (1.2 mL). After 5, 10, 15, 30, 45, 60, 90, 120, 180, and 240 min, 600 μL of medium was extracted from the basolateral compartment and replaced by the same volume of fresh transport medium to maintain the total volume in the chamber constant. The fluorescence in the extracted medium, which is a measure of the polymeric micelles that crossed the epithelium monolayer, was quantified by fluorescence spectrophotometry in a Fluoroskan Ascent Plate Reader (Thermo Fisher Scientific Oy) using black 96-well flat bottom plates (Greiner BioOne, Kremsmunster, Austria) at wavelengths of 485 nm for excitation and 538 nm for emission and the results interpolated in a calibration curve built with different polymeric micelles concentrations (in the 0.0001−0.1% w/v range) containing of FITC-labeled 25% w/w CBD-loaded polymeric micelle concentrations (R^2^ = 0.999). The apparent permeability (Papp) was calculated according to Equation 2

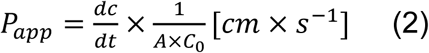

where *dc/dt* is the permeability rate of the polymeric micelles (expressed in µg/s) across the monolayer, C_0_ is the initial concentration of the polymeric micelles in the donor compartment (expressed in µg/cm^3^), and A is the surface area of the membrane (1.131 cm^2^). Results were expressed as mean ± S.D. of three independent experiments.

### 2.4. Pharmacokinetic and pharmacodynamic studies

All experiments involving Sprague-Dawley rats were conducted in accordance with the ethical standards and guidelines set by the Institutional Animal Research Ethical Committee. In the *in vivo* experiments, we addressed ASD-like symptoms, focusing on the social cooperation behavior.^34^

#### 2.4.1. Accumulation of CBD-loaded polymeric micelles in the olfactory bulb and the brain of autistic rats upon intranasal administration

To assess the brain accumulation of 25% w/w CBD-loaded Pluronic^®^ F127 polymeric micelles upon i.n. administration, the copolymer was fluorescently labeled by the conjugation of near-infrared fluorescent dye NIR-797 (Sigma-Aldrich). In brief, Pluronic^®^ F127 (50 mg) was dissolved in MilliQ-water (1 mL) with pH of 5.5. Then, NIR-797 (0.4 mg) was dissolved in DMSO (500 µL) and added to the copolymer solution. The reaction was allowed to proceed under magnetic stirring (300 RPM) for 12-16 h, at RT and protected from light. After the reaction, the resulting solution was dialyzed against Milli-Q water for three days using regenerated cellulose dialysis membranes (molecular weight cut-off of 3500 Da) and freeze-dried (Labconco Free Zone 4.5 plus L Benchtop Freeze Dry System). To synthesize NIR-797-labeled polymeric micelles, 20% w/w of F127-NIR was mixed with 80% w/w of unlabeled Pluronic^®^ F127. Then, NIR-797-labeled 25% w/w CBD-loaded polymeric micelles were prepared as described above for FITC-labeled ones. To assess the accumulation in the brain, NIR-797-labeled 25% w/w CBD-loaded polymeric micelles were administered i.n. to rats (CBD dose of 5 mg/kg; volume of 0.205 µL/g). For example, a 200-g rat was administered 42 µL of the micellar suspension, the total volume being administered in two rounds: 10 µL per nostril in the first round and 11 µL per nostril in the second one. All the samples used in the experiments were freshly prepared. To track the accumulation of the CBD-loaded Pluronic^®^ F127 polymeric micelles in the brain, animals were sacrificed by decapitation after 20, 45, 90 and 180 min, and their brains were carefully dissected. We also dissected olfactory bulbs 90 min after administration. Imaging was performed using an IVIS^®^ Spectrum CT system from PerkinElmer (Waltham, MA, USA). The imaging was carried out using filters that corresponded to the excitation (780 nm) and emission (831 nm) peaks of the NIR-797 dye. The radiant efficiency was quantified using a region-of-interest (ROI) in the acquired images. This quantification was facilitated by the Living Imaging analysis software (PerkinElmer).

#### 2.4.2. CBD pharmacokinetics in autistic rats

The CBD concentration in the brain and plasma of autistic rats over time was studied after (i) i.n. administration of 25% w/w CBD-loaded Pluronic^®^ F127 polymeric micelles (CBD dose of 5 mg/kg; volume of 0.205 µL/g), (ii) i.n. administration of 2.43% w/v CBD solution in propylene glycol (CBD dose of 5 mg/kg; volume of 0.205 µL/g) and (iii) oral administration of 25% w/w CBD-loaded Pluronic^®^ F127 polymeric micelles (CBD dose of 15 mg/kg; volume of 0.615 µL/g). For this, male ASD rats (300-360 g) were randomly divided into equal groups for testing the three treatments at five time points. Blood samples (0.5-1 mL) from cardiac puncture were taken at 20, 45, 60, 120 min and 24 h after administration, animals sacrificed by decapitation and the brains carefully collected. Then, CBD was quantified in plasma and brain tissue by high performance liquid chromatography (HPLC) according to a previously a described method with some modifications.^38^ Briefly, plasma samples were centrifuged at 3000 RPM for 10 min at 4°C (Hermle Labortechnik GmbH, Wehingen, Germany) and brains were homogenized by ultrasonication (ultrasonic processor, Cole-Parmer, Vernon Hills, IL, USA) with 1.5 mL/g phosphate buffered saline (PBS, Sigma-Aldrich) During the homogenization process, the vials containing the brains and PBS were kept on ice. CBD was extracted from plasma and brain homogenates (100 µL) by adding a hexane:ethyl acetate:chloroform co-solvent mixture (all of them supplied by Sigma-Aldrich) with a 4:1:1 volume ratio (220 µL), vortexing for a few minutes, and centrifuging for 20 min at 13,500 RPM (Hermle Labortechnik GmbH, Wehingen, Germany). After centrifugation, the organic supernatant was collected and dried under nitrogen flow. Dry residues were reconstituted with an acetonitrile:methanol mixture (supplied by Avantor, Inc., Radnor Township, PA, USA, respectively) with 1:1 volume ratio (100 µL). Samples were filtered (PTFE CR Acrodisc Syringe Filter, 13 mm, 0.45 µm, Waters Corp., Milford, MA, USA), and injected into a high-performance liquid chromatograph (HPLC, Alliance HPLC System, Waters Corp.) and CBD quantified with a UV-Vis detector at λ = 228 nm with mobile phase of acetonitrile:Milli-Q water (65:35 volume ratio), respectively, using a C18 column with flow rate of 0.8 mL/min and run time of 35 min. The concentration from the HPLC elution peaks was calculated using a calibration curve of different concentrations of CBD in rat plasma and brain homogenates with concentrations between 28 and 3600 ng/mL (R^2^ = 0.990 for plasma and R^2^ = 0.997 for brain). The pharmacokinetic (PK) parameters C_max_ which is the maximum concentration of CBD, t_max_ which is the time to reach the C_max_, the t_1/2_ which is the half-life, and the area-under-the-curve (AUC_0-24 h_) which is a measure of the bioavailability, were calculated using the PKSolver add-in software.^39^

#### 2.4.3. Pharmacodynamic experiments – Behavioral tests in rats

Behavioral tests that are characteristic of ASD were conducted in this rat ASD model. Three samples were systematically compared: (i) i.n. administration of 25% w/w CBD-loaded Pluronic^®^ F127 polymeric micelles (CBD dose of 5 mg/kg; volume of 0.205 µL/g); (ii) i.n. administration of unloaded 10% w/w Pluronic^®^ F127 polymeric micelles (volume of 0.205 µL/g) and (iii) oral administration of 25% w/w CBD-loaded Pluronic^®^ F127 polymeric micelles (CBD dose of 15 mg/kg; volume of 0.615 µL/g).

*Social cooperation:* The SC learning is carried out in a dimly lit room (8lux) using a custom-made black lusterless Perspex box (40 cm W, 120 cm L, 50 cm H) as we previously described,^33^ divided vertically into two lanes by transparent-perforated divider, allowing visual, auditory, tactile, and olfactory interactions between the two rats. The definition of the maze is based on a virtual three zones that perpendicularly divide the two lanes (A = start section, B = middle and C = end). When the two rats coordinate their movement throughout the three virtual zones (A to C), and the trial’s requirements are met by the algorithm programmed in the Ethovision XT 11.5 software (Noldus, Wageningen, The Netherlands), both rats receive 70 μL of 20% w/v sucrose solution in water as reinforcement. During the pre-learning phase, rats are habituated individually to the cooperation maze for a single session of 10 min and are allowed to consume sucrose at the end of the maze to overcome neophobia. Throughout the cooperative learning period, rats are water limited for 16–18 h a day before the daily trial. The aim of limiting the availability of water is to enhance the motivation of the rats to perform the task. On the first day of learning and onward (for 18 consecutive days), each rat is placed in the maze with the same partner (coming from a different home cage) for a 15-min session. A trial is considered successful when the rats shuttle coordinately between the three predefined zones (“A” to “B” to “C”) within a maximal 10 s delay from the adjacent zone and arrive at the end of the maze together (zone “C”). An iterated trial starts when both rats return to zone “A”. The rats do not receive a reward at the end of zone “C” if they are in different zones for more than 10 s or if they are more than one zone apart (“A” and “C”). During the first 12 days, the rats are required to cross from area A to area B in order to receive a reward, from day 13 to 18, they had to cross from area A to B and then to C consecutively to receive the reward (**Figure S1A**). The behavior is monitored and recorded by auEye UI-308x-CP-C camera (IDS Imaging Development Systems GmbH, Obersulm, Germany) and analyzed by a custom-made algorithm using Ethovision XT 11.5 software.^34^ The rats received the treatments once a day, 20 min before entering the maze, along 18 days of learning.

*Open field – Anxiety and activity measurements:* The open field apparatus is made of black lusterless Perspex box (100L × 100W × 40H cm) placed in a dimly lit room (50 lux). Rats are placed in the corner of the open field (facing the wall) and given 5 min of free exploration. The behavior is monitored by a uEye UI-308x-CP-C camera and analyzed with Ethovision XT 11.5 software, as we previously described (**Supplementary Figure S1B**).^40^

*Object recognition – Selective attention:* The test is conducted in a black lusterless Perspex box (100L × 100W × 40H cm) placed in a dimly lit room (50 lux). Rats are acclimated to the test arena for 10 min on the first day (i.e., they are allowed to explore the arena without any objects). Initially, rats are placed in the distal part of the box, facing the wall. On the second day, rats are allowed to explore the arena for 10 min, with the presence of four identical objects (Lego cubes). Next, rats are removed to their home cages for 1 h, then placed back in the arena (which was cleaned) for a 5 min test trial with 3 familiar and identical objects, and one new object (that is different in size, shape, and color). The novel preference (%) was calculated as the proportion of time spent exploring the novel object relative to the familiar objects were monitored by a uEye UI-308x-CP-C camera and analyzed with Ethovision XT 11.5 software, as previously described (**Supplementary Figure S1C**).^40,41^

*Water maze – Spatial learning:* The water maze is a circular tank, filled with water (1.4 m in diameter with a rim 0.6 m high) made from black plastic. Water depth is 45.0 cm, and temperature is maintained at 23°C ± 1. The only mean of escape from the water is a 10 cm in diameter hidden platform, whose top lies 1 cm beneath the water surface. The water maze itself has no landmarks; the only landmarks are outside the water maze, on the surrounding walls. Training consisted of four trials per day, with a 3-min inter-trial interval, for 4 consecutive days. Rats were randomly placed (facing the wall) at one of four starting positions (N, E, W, and S) and allowed to swim freely for a maximum of 90 s or until reaching the hidden platform. At the end of each trial, each rat was led to or left on the platform for 15 s. The behavior is monitored by a uEye UI-308x-CP-C camera and analyzed with Ethovision XT 11.5 software, as we previously described (**Supplementary Figure S1D**).^41^

*Social ultrasonic vocalization:* Rats are placed in a social interaction box (100 X 50 cm) without any divider, enabling free social interaction and communication for 5min. The box is placed in a sound-attenuating room. Vocalizations are recorded from the interacting animals, by placing a high-frequency sensible microphone (Motu, MA, USA) 2 cm from the bottom of the box facing up 20° from the floor and covered by a metal mesh. Ultrasonic vocalizations between 30-70kHz are counted (**Supplementary Figure S1E**).

### 2.5. Data and statistical analyses

All statistical analyses are conducted using GraphPad Prism software version 7.04 (GraphPad Software, San Diego, CA, USA). Results are reported as the mean ± S.D or mean ± S.E. (for PK studies) of at least three independent experiments. A *p*-value of at least 5% is considered significant for all tests.

## 3. Results and Discussion

### 3.1. Preparation and characterization of CBD-loaded polymeric micelles

The physicochemical properties and hepatic first-pass metabolism reduce the oral bioavailability and stability of CBD in the biological milieu.^12,42^ We hypothesized that the i.n. administration of CBD-loaded nanocarriers would localize the delivery,^20,43^ reducing the required dose to achieve a therapeutic effect and systemic exposure and the associated off-target toxicity.^44^ Polymeric micelles are among the most versatile nanocarriers for delivery and targeting of lipophilic cargos in general^45,46^ and by mucosal routes in particular.^47,48^ With a translational vision, we decided to utilize a polymeric amphiphile that is biocompatible, enables the preparation of the polymeric micelles by a simple and scalable method, it is approved by the US Food and Drug Administration as pharmaceutical and it is commercially available in pharmaceutical grade, namely Pluronic^®^ F127.^49^

To optimize the nanoformulation, we systematically evaluated the encapsulation capacity of 10% w/w Pluronic^®^ F127 polymeric micelles in water by gradually increasing the CBD amount to be encapsulated by the simple diffusion method.^48^ The encapsulation limit was 27% w/w. Even if transparent to the naked eye (data not shown), 27% w/w CBD-loaded polymeric micelles were physically unstable, and CBD precipitated over time. Systems with a higher CBD loading of 30% w/w showed relatively fast precipitation of part of the payload. Thus, we continued our work with a 25% w/w CBD-loaded nanoformulation that was transparent to the naked eye, homogeneous and physically stable (**Figure 1A**) with a monomodal size population, a D_h_ by intensity of 41 ± 1 nm (and by number of 23 ± 1 nm), and a low PDI of 0.277 ± 0.020, as measured by DLS (**Figure 1B**). The size, the concentration, and the Brownian motion of the polymeric micelles in suspension were characterized using NTA (**Supplementary Video S1**). The D_h_ by NTA (40 ± 5 nm) was similar the one measured by DLS (**Figure 1C**) and the concentration 3 x 10^9^ nanoparticles/mL (after correction by the dilution factor). The Z-potential is an estimation of the surface charge, and it plays a fundamental role in the interaction of nanoparticulate matter with mucus and cells. PEO-PPO block copolymers are non-ionic, though their polymeric micelles usually display a slightly negative Z-potential that depending on the hydrophilic-lipophilic balance of the copolymer ranges between −7 and −14 mV,^50^ and has been reported to improve nanoparticle retention in mucosal tissues.^51^ CBD-loaded Pluronic^®^ F127 polymeric micelles showed a value −10 ± 2 mV that is in very good agreement with the literature.^52^

**Figure 1.**
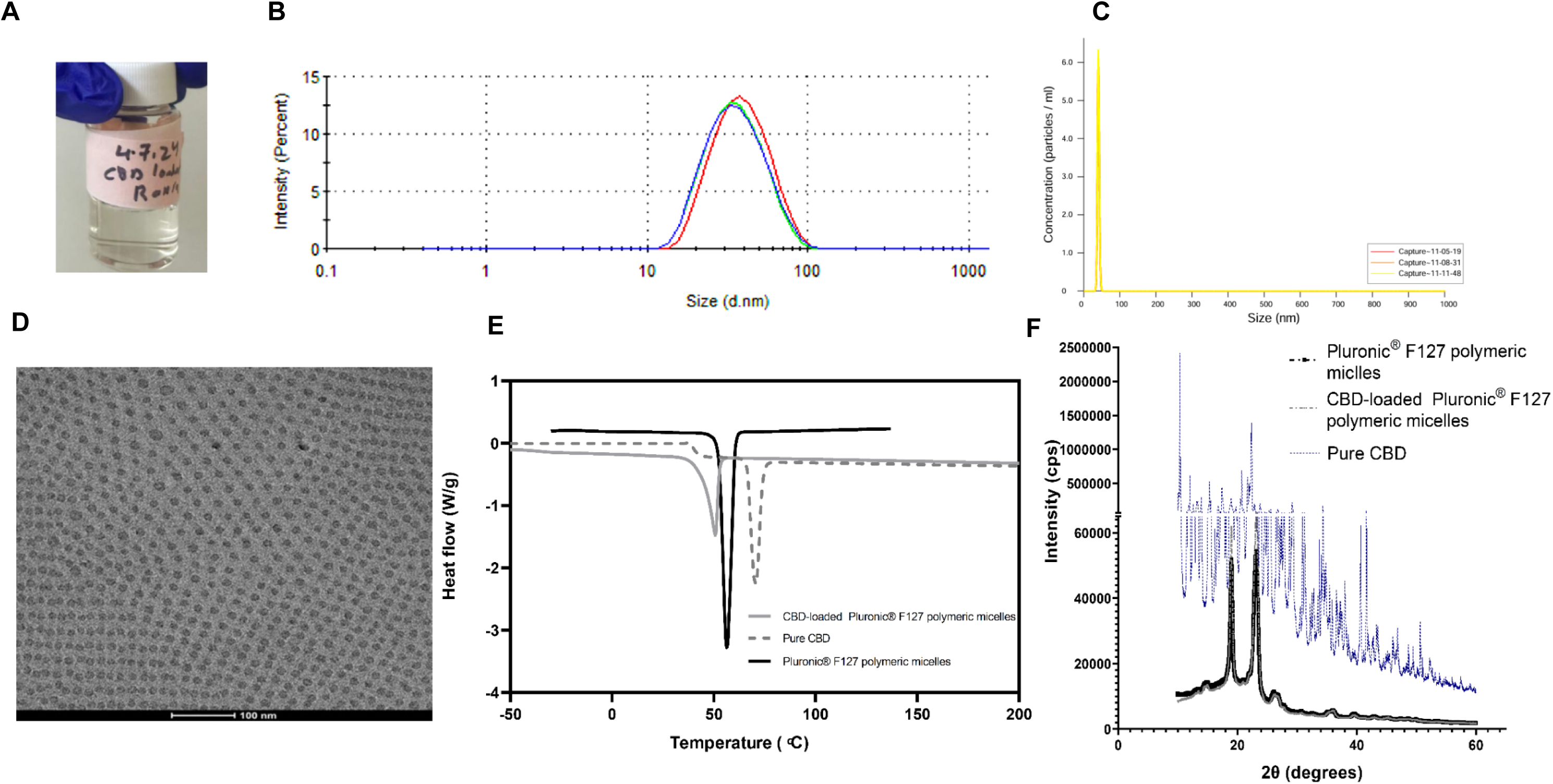
Characterization of CBD-loaded Pluronic^®^ F127 polymeric micelles. (A) A nanoformulation of 25% w/w CBD-loaded Pluronic^®^ F127 polymeric micelles. (B) Size distribution (expressed as D_h_ by Intensity), as measured by DLS at 25°C. (C) Size distribution, as measured by NTA at 37°C. (D) Representative cryo-TEM micrograph. (E) Thermal properties of pure CBD, and unloaded and CBD-loaded polymeric micelles, as characterized by DSC. (F) Diffractogram of pure CBD, and unloaded and CBD-loaded polymeric micelles, as characterized by PXRD.

To gain insight into the morphology and nanostructure of the CBD-loaded Pluronic^®^ F127 polymeric micelles, we analyzed them by cryo-TEM. Void PEO-PPO polymeric micelles are hardly visualized by this technique as dark dots due to the limited difference in electronic density and contrast between the two copolymer blocks, PEO and PPO, and some modifications were investigated to enable a more detailed characterization of the nanostructure.^53^ CBD nanoencapsulation substantially increased the contrast between the hydrophilic PEO shell and the hydrophobic PPO core, revealing a spherical shape and the presence of the two well-define micellar domains, the core and the shell (**Figure 1D**). It also confirmed the absence of free CBD outside the polymeric micelles.

Lipophilic cargos encapsulated within polymeric nanoparticles are often dispersed at the molecular level within the polymeric matrix and in amorphous form, this phenomenon being more likely in self-assembled systems. The release of amorphous cargos from the polymeric matrix is often faster than that of crystalline ones. To investigate the thermal properties of CBD within the polymeric micelles, pure CBD, and unloaded and CBD-loaded Pluronic^®^ F127 polymeric micelles, were analyzed by DSC. Pure CBD displayed a sharp melting endotherm at 70°C with a melting enthalpy (ΔH_m_) of 68.15 J/g) due to its crystalline nature,^54^ while unloaded polymeric micelles the melting temperature (T_m_) of PEO blocks at 55°C (ΔH_m_ = 109.55 J/g) (**Figure 1E**); PPO is fully amorphous. The CBD-loaded nanoformulation exhibited a single endothermic peak at 50°C (ΔH_m_ = 79.10 J/g) that matched the melting point of PEO, and the complete absence of any transition assignable to crystalline CBD (**Figure 1E**). A decrease in the T_m_ and ΔH_m_ of PEO in the loaded polymeric micelles indicates that CBD hinders its crystallization process. These results indicate the amorphous nature of CBD within the polymeric micelles.

To complement the analysis, the same samples were analyzed by PXRD. The diffractogram of pure CBD showed distinct sharp peaks, consistent with its crystalline structure (**Figure 1F**). Unloaded Pluronic^®^ F127 polymeric micelles exhibited two characteristic diffraction peaks at 2θ ≈ 19° and 23°, corresponding to crystalline PEO segments. CBD-loaded polymeric micelles presented only the two peaks of crystalline PEO, confirming the CBD amorphization upon encapsulation.

Formulations for nasal administration need to comply with low viscosity to enable the easy instillation and pH in the 4.5-6.5 range to minimize irritation of the nasal mucosa and epithelium. Our nanoformulation exhibited a pH of 6.2 ± 0.1 and a low dynamic viscosity of 6.0 ± 0.5 mPa.s which are compliant with this administration strategy.

### 3.2. CBD release in vitro

The release profile of CBD from Pluronic^®^ F127 polymeric micelles displayed a typical 2-stage profile, with a burst release of ∼40% during the first 3 h of the assay and a more gradual one that reached 75% and 85% after 10 and 24 h, respectively (**Supplementary Figure S2**). To understand the release mechanism, the cumulative release data were fitted to different release models using DDSolver add-in software.^36^ The Peppas–Sahlin model provided the best fit (R^2^ = 0.98). This model is described by Equation 3

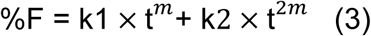

where %F is the cumulative drug release percentage at time t, k1 is the rate of drug release due to diffusion, k2 is the rate of drug release due to polymer relaxation or erosion, and m is a diffusional exponent.

The first term (k1 × t^𝒎^) reflects Fickian diffusion, while the second one (k2 × t^2𝑚^) accounts for polymer relaxation or erosion mechanisms. In our case, an *m* value of 0.671 indicated that both mechanisms contribute to the release of CBD from the polymeric micellar system.^55^

### 3.3. Cell compatibility and permeability across model of the nasal epithelium in vitro

Towards the PK and PD (behavioral) studies in a preclinical model of ASD, we studied the cell compatibility and permeability of 25% CBD-loaded Pluronic^®^ F127 polymeric micelles in the only commercially available human nasal epithelium cell line, RPMI 2650. PEO-PPO block copolymers have demonstrated excellent cell and biocompatibility by various administration routes.^31^ Conversely, CBD assays in a plethora of cell types indicated that concentrations above ∼2-10 µM are cytotoxic.^56^ Ensuring low cell toxicity was crucial to assess the permeability of this nanoformulation in RPMI 2650 monolayers without affecting its integrity and confluence due to cell death. For this, 25% w/w CBD-loaded polymeric micelles were diluted to different extents in the cell culture medium to achieve final CBD concentrations between 1.178 and 8.588 µM (equivalent to micellar concentrations of 0.005% and 0.25% w/v, respectively), exposed to the loaded polymeric micelles for 24 h and the viability estimated by the MTT assay. In good agreement with the literature, high cell viability >89% was recorded for a CBD concentration of up to 5.732 µM. A more concentrated sample led to a viability loss of 40% *in vitro* (**Figure 2A**). In advance, the permeability studies *in vitro* were conducted, CBD-loaded Pluronic^®^ F127 polymeric micelles showed good permeability under ALI conditions which increase the formation of epithelial tight junctions, 0.01% and 0.05% w/v polymeric micelles showing Papp values of 55.43 ± 5.89 and 32.50 ± 6.46 cm/s × 10^-7^, respectively (p < 0.05) (**Figure 3B**). A decrease in the Papp of a more concentrated micellar system probably stemmed from some level of saturation of the permeability pathways. These results correlate with good nasal permeability *in vivo*.^57^

**Figure 2.**
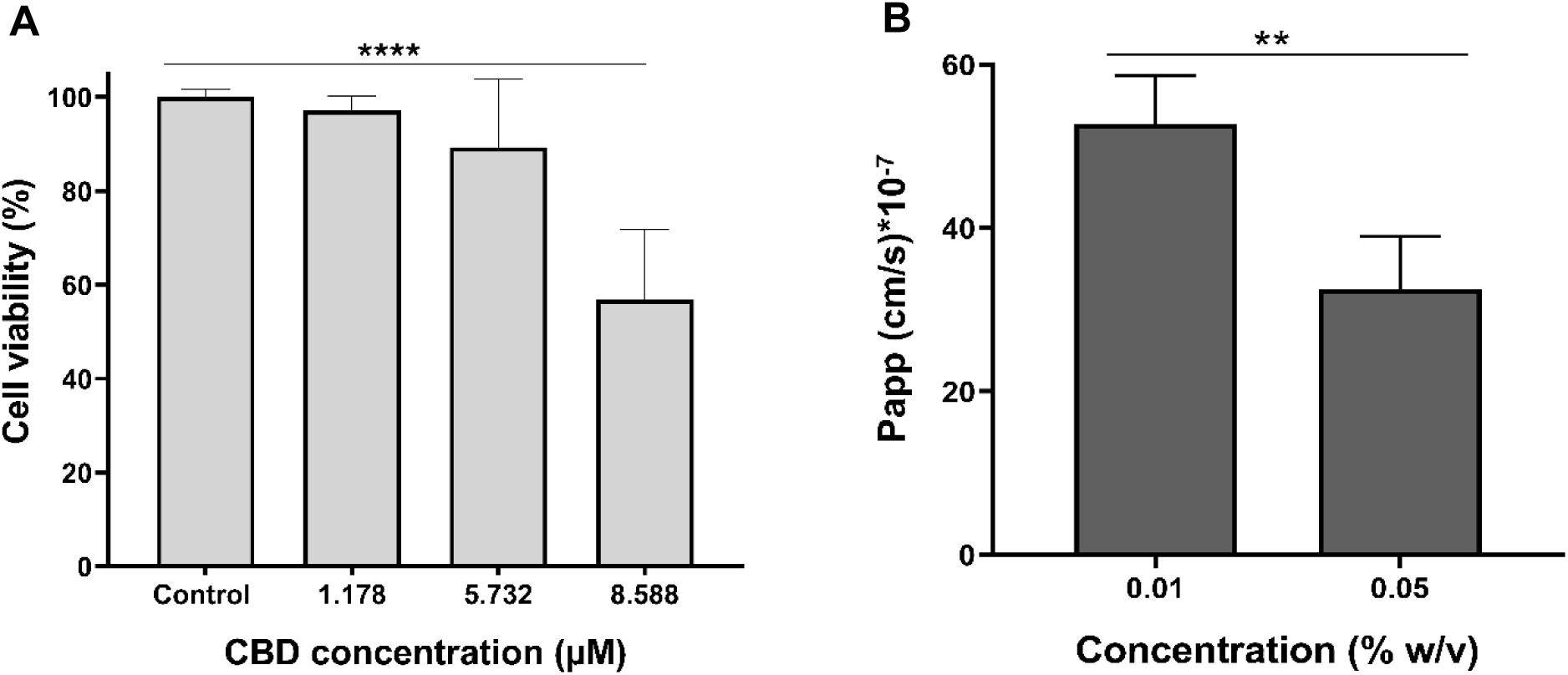
Compatibility and permeability studies of CBD-loaded Pluronic^®^ F127 polymeric micelles in the human nasal epithelium cell line RPMI 2650. (A) Cell viability upon exposure to micellar systems with different final CBD concentrations for 24 h at 37°C, as estimated by the MTT assay (n = 3). The original 25% w/w CBD-loaded Pluronic^®^ F127 polymeric micelles were diluted in culture medium to final concentrations of 0.005-0.25 % w/v. All data are presented as mean ± S.D. respectively (p < 0.0001). (B) Apparent permeability coefficient (Papp) of 0.01% and 0.05% w/v CBD-loaded Pluronic^®^ F127 polymeric micelles under ALI conditions (n = 6). ** Statistically significant difference (p < 0.01) and **** statistically significant difference (p < 0.0001).

**Figure 3.**
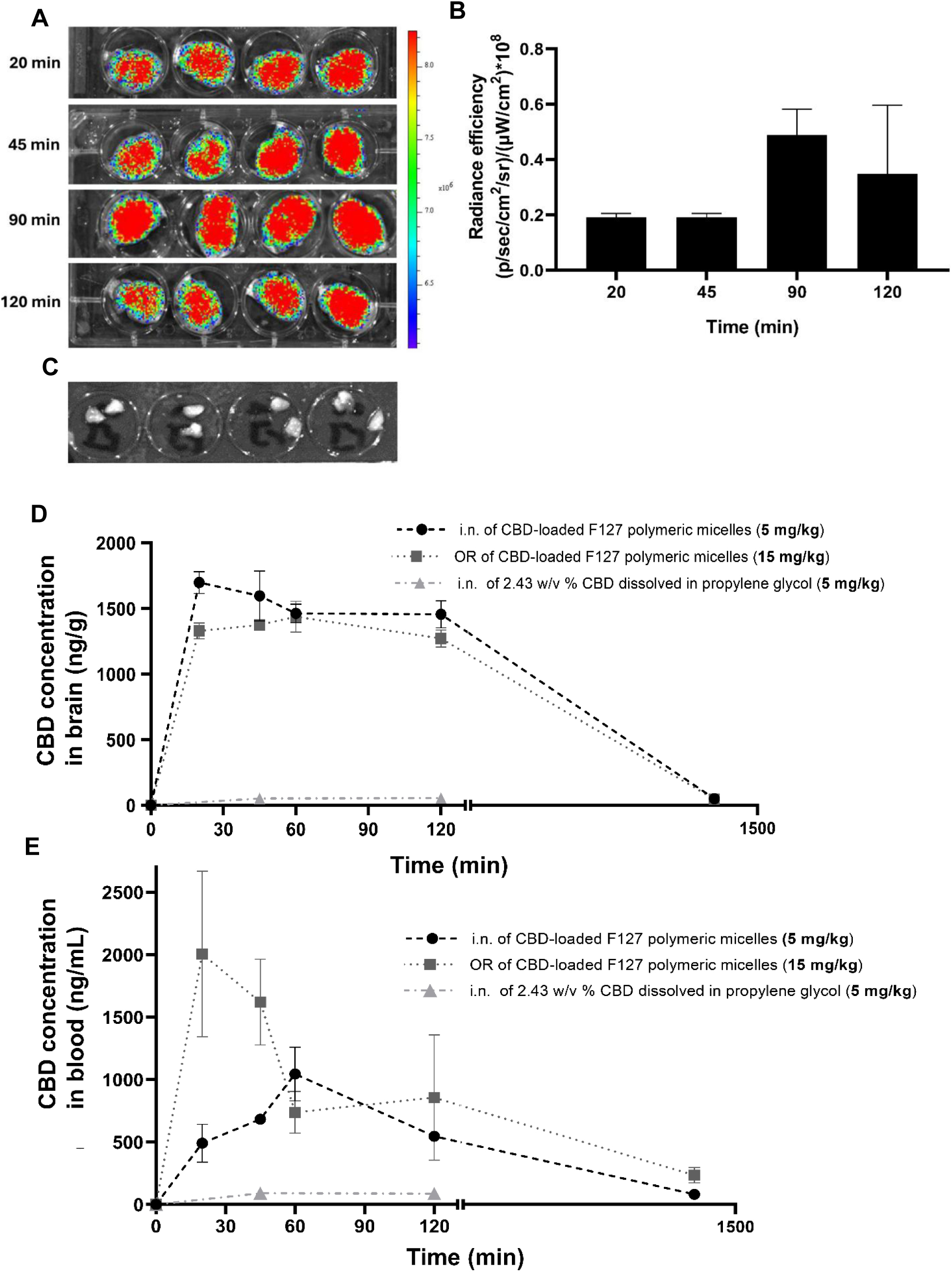
Pharmacokinetics of 25% w/w CBD-loaded Pluronic^®^ F127 polymeric micelles in autistic rats. (A) IVIS fluorescence micrographs of rat brains at different time points after i.n. administration of NIR-797-labeled CBD-loaded polymeric micelles, as imaged by IVIS (n = 4). (B) Fluorescence radiance intensity profiles in the brain of rat (n = 4). (C) IVIS fluorescence micrographs of rat olfactory bulbs 90 min after i.n. administration of NIR-797-labeled CBD-loaded polymeric micelles, as imaged by IVIS (n = 4). (D) CBD concentration in rat brains following the i.n. and oral administration of 25% w/w CBD-loaded Pluronic^®^ F127 polymeric micelles and i.n. administration of CBD dissolved in propylene glycol. (E) CBD concentration in rat plasma following the i.n. and oral administration of 25% w/w CBD-loaded Pluronic^®^ F127 polymeric micelles and i.n. administration of CBD dissolved in propylene glycol. In all the PK studies, the i.n. and oral CBD dose was 5 and 15 mg/kg, respectively. All data are presented as mean ± S.E. (n = 4).

### 3.3. Pharmacokinetic and pharmacodynamic studies

#### 3.3.1. Accumulation of CBD-loaded polymeric micelles in the olfactory bulb and the brain of autistic rats upon intranasal administration

Our therapeutic strategy relies on the nose-to-brain transport of the polymeric micelles for which they need to initially cross the nasal mucus layer and epithelium, their accumulation in the brain and the local release of the cargo.^58^ To assess the capacity of the CBD-loaded 25% w/w Pluronic^®^ F127 polymeric micelles to accumulate in the brain of autistic rats, we labeled them by the conjugation of the fluorescent dye NIR-797 to the terminal hydroxyl moieties of the block copolymer, administered them intranasally (CBD dose of 5 mg/kg; volume of 0.205 µL/g) and tracked their accumulation in the olfactory bulb and the brain at different time points by IVIS bioimaging. The loaded polymeric micelles could be detected in the brain of the rats even 20 min post-administration and for at least 2 h (**Figure 3A**), with a maximum fluorescence recorded after 90 min (**Figure 3B**); the background fluorescence of the brains of untreated rats was used as a blank.

Intranasally administered nanoparticles can be transported to the brain along the olfactory bulb.^59,60^ To evaluate whether the polymeric micelles are retained in this region of the CNS, we dissected the olfactory bulbs 90 min post-administration and imaged them. No fluorescence could be detected, in line with the successful transport to the brain (**Figure 3C**).

#### 3.4.2. CBD pharmacokinetics in autistic rats

Once we confirmed that the polymeric micelles accumulate in the brain of autistic rats, we studied the PK of CBD in the brain and plasma after the i.n. administration of the loaded polymeric micelles (CBD dose of 5 mg/kg; volume of 0.205 µL/g) and compared the PK parameters with those of the same nanoformulation administered orally (CBD dose of 15 mg/kg; volume of 0.615 µL/g) and the i.n. administration of a 2.43% w/v CBD solution in propylene glycol (CBD dose of 5 mg/kg; volume of 0.205 µL/g). The oral route is the most popular for CBD administration in children, while propylene glycol is often used in PK and PD studies as a vehicle for poorly water-soluble drugs.

In the brain, the C_max_ following i.n. administration of CBD-loaded Pluronic^®^ F127 polymeric micelles was 1699 ± 44 ng/g and significantly higher than that of the oral administration (C_max_ = 1437 ± 53 ng/g, p < 0.01), despite the 3-fold smaller i.n. dose (**Figure 3D**). These results demonstrate the advantage of the i.n. route over the oral one. In addition, the t_max_ was shorter (20 ± 21 min versus 60 ± 19 min) which may have key implications in the onset of pharmacological activity (**Table 1**). Despite the 3-fold decrease in the CBD dose, in the brain, the AUC_0-24 h_ for i.n. administration (1163 ± 86 µg/ng x min) was significantly greater than that of the oral administration (1020 ± 80 µg/ng x min, p < 0.01) (**Table 1**). No differences were observed in the t_1/2_. In plasma, i.n. administration led to a statistically significant decrease in the C_max._ The plasma C_max_ following i.n. administration was 1045 ± 215 ng/mL, detected at 60 min post-administration, compared to 2126 ± 594 for oral administration: t_max_ values being 20 ± 10 and 60 ± 20 min for the oral and i.n. routes, respectively (**Figure 3E**, **Table 1**). This difference was expected because i.n. administration primarily targets the brain, limiting the drug fraction entering the systemic circulation and highlights the advantage of this alternative administration route to minimize systemic exposure and the potential off-target toxicity. The lower systemic exposure was confirmed by a sharp, though not statistically significant decrease, in the AUC_0-24 h_ from 850 ± 393 to 494 ± 19 µg/mL x min for i.n. and oral routes, respectively (**Table 1**).

**Table 1.** Pharmacokinetic parameters of CBD in autistic rat plasma and brain after the i.n. and oral administration of 25% w/w CBD-loaded Pluronic^®^ F127 polymeric micelles. The i.n. and oral dose was 5 and 15 mg/kg, respectively. All data are presented as mean ± S.E. (n = 4).

| Administration route | PK parameter |  |  |  |  |  |  |  |
| --- | --- | --- | --- | --- | --- | --- | --- | --- |
|  | Brain |  |  |  | Plasma |  |  |  |
| | $t_{\max} \pm \text{S.E.}$<br>(min) | $C_{\max} \pm \text{S.E.}$<br>(ng/g) | $t_{1/2} \pm \text{S.E.}$<br>(min) | $AUC_{0-24h}$<br>($\mu\text{g/g} \times \text{min}$) | $t_{\max} \pm \text{S.E.}$<br>(min) | $C_{\max} \pm \text{S.E.}$<br>(ng/mL) | $t_{1/2} \pm \text{S.E.}$<br>(min) | $AUC_{0-24h}$<br>($\mu\text{g/mL} \times \text{min}$) |
| Intranasal | 20 ± 21 | 1699 ± 44 ** | 279 ± 10 | 1163 ± 86 ** | 60 ± 20 * | 1045 ± 215 | 417 ± 22 | 494 ± 19 |
| Oral | 60 ± 19 | 1437 ± 53 | 278 ± 3 | 1020 ± 80 | 20 ± 10 | 2126 ± 594 | 480 ± 21 | 850 ± 393 |

\* Statistically significant difference between i.n. and oral routes ( $p < 0.05$ ).
\*\* Statistically significant difference between i.n. and oral routes ( $p < 0.01$ ).

The longer t_max_ in plasma for i.n. administration indicates that after reaching the brain, CBD is redistributed to the systemic circulation from which it can be cleared from the body, as indicated by the very similar t_1/2_ values. Additionally, we evaluated the concentration of CBD in the brain and blood following i.n. administration of CBD dissolved in propylene glycol, without nanocarriers (CBD dose of 5 mg/kg). The results demonstrated that CBD reached the brain and the blood at very low levels, with concentrations of 52 ± 6 ng/g in the brain and 87 ± 8 ng/mL in the blood, after 45 min. Overall, our PK results confirm the advantage of i.n. over oral CBD delivery and support the conduction of PD (behavioral) studies in autistic rats. It is also remarkable that the polymeric micelles led to a dramatic increase in the oral bioavailability of CBD when compared to simple solutions,^61^ which may pave the way for its implementation in systemic pharmacological interventions.

#### 3.3.3. Pharmacodynamic experiments – Behavioral tests

Though ASD is increasingly prevalent,^62^ with a 3:1 male:female ratio,^1^ there is no designated pharmacological treatment specifically developed for ASD. Therefore, psychotropic medications are commonly prescribed, although frequently without sufficient effect and with bothersome side effects that are especially challenging in the pediatric population.^63^ Our goal was to develop and preclinically assess the efficacy of a new scalable nanoformulation of CBD for intranasal delivery to the brain and examine its effects on ASD-like behaviors. The exist different translational rodent models recapitulating some of the core behavioral symptoms of autism that have been utilized to screen pharmacological interventions.^64,65^ However, there is an enormous need to develop revised theoretical approaches accompanied by new animal model research methodologies and tools, to revolutionize the understanding and treatment of ASD. The main theoretical approach to investigating ASD is ‘bottom-up’, in which clinically relevant genes are identified in human individuals with ASD and nurturing specific genetic mutations in animal models.

This strategy has not led to designated treatments.^66,67^ Despite the recognition of nearly 800 candidate genes for ASD, no designated pharmacological treatments for ASD have been identified.^65,66^ Though some promising genetic animal models have been investigated, the attrition rate of translating these models to human studies is very high and of limited clinical impact.^68,69^

In this work, we implemented a ‘top-down’ approach by focusing on the transgenerational inheritance of social cooperation (SC), while paying attention to functioning heterogeneity. SC behavior is a widespread phenomenon in pairs or large groups aimed to achieve a tangible, immediate reward, by voluntary joint action.^70^ Cooperative behavior, in animals and humans, is a complex executive function, requiring various sensory, emotional, cognitive and social skills, and necessitating concurrent monitoring of ongoing social relationships,^71^ by pronounced abilities of social decision-making and reinforcement learning.^72,73^ Thus, SC behavior seems to depict the core symptom of ASD and reflects its complexity. Thus, utilizing our computerized SC maze,^34^ a pair of rats are required to manifest similar executive function abilities as in socially interacting humans. An important and unique characteristic of the SC maze is the exclusion of physical touch, which is a part of previously introduced SC in other animals.^74,75^ Excluding physical touch as a mean of social communication between the cooperating rats may exacerbate alternative, non-physical communication, i.e. vocalizationThis ability provides not just adequate sensory communication between the rats, but rather mutual understanding of how to perform the next step towards the reward. The heritability estimate for ASD varies between 50% and 90%,^76,77^ however, its molecular diagnostic yield is extremely low. Relating to the inheritance factors, a selective breeding procedure was conducted along 15 generations, in which the selection rule was based on defining the highest or lowest 10% of SC performers, yielding two distinct subpopulations of ’High’ and ’Low’ SC that progressively differed in SC performance, with the Low SC presenting an ASD-like behavioral phenotype (data not shown).

Low-performance female and male rats (weight range of 200-330 g) were divided into four groups, as follows (i) 5 mg/kg of i.n. administration of 25% w/w CBD-loaded Pluronic^®^ F127 polymeric micelles (volume of 0.205 µL/g) (i.n.+, n = 36, 22 female and 14 male), (ii) 15 mg/kg of oral administration of 25% w/w CBD-loaded Pluronic^®^ F127 polymeric micelles (OR+, n = 36, 22 female and 14 male), (iii) i.n. administration of the equivalent volume (0.205 µL/g) of unloaded Pluronic^®^ F127 polymeric micelles (i.n-, n = 18, 10 female and 8 male) and (iv) untreated rats (Low, n = 32, 16 female and 16 male). The formulations were administered every 24 h (once-a-day, 18 administrations) and the SC test was carried out every day 20 min post-administration for 18 days. Results revealed statistically significant differences in cooperative behavior among the i.n.+ group and the rest over time (**Figure 4A**). The i.n.+ group demonstrated a sharp improvement in cooperation (the number of recorded rewards increased from 1-2 at the beginning of the test to ∼10 at day 18), differences being statistically significant with respect to the untreated group from day 7 (except day 8) until day 18 (p < 0.001) and the OR+ group on days 7 (p < 0.01), 9-12 (p < 0.001), 13 (p < 0.01), 14 (p < 0.001), 17 (p < 0.001) and 18 (p < 0.001). In addition, the i.n.+ group displayed significantly higher cooperation on most days than the i.n.-group: days 7 (p < 0.05), 9-12 (p < 0.001), 13 (p < 0.05), 14 (p < 0.001), 15-16 (p < 0.05) and 17-18 (p < 0.001) (**Figure 4A**). The OR+ group showed a more moderate increase in cooperation from 1-2 to 5 rewards at the end of the treatment, even though the oral dose was 3-fold higher than the i.n. one (**Figure 4A**); these reward numbers being statistically significant in few days with respect to the control rats in days 11 (p < 0.05), 14 (p < 0.01), 15 (p < 0.05), 16 (p < 0.01), and those treated with unloaded polymeric micelles (i.n.-) only in days 16 (p < 0.01) and 18 (p < 0.01). The transient drop in cooperation observed on day 13 across all the groups could be attributed to the increased difficulty of the test from this time point on. Between days 0 and 12, the rats need to cross from section A to section B to get the reward, while from day 13 on, they are required to cooperate and cross three zones (A to B and to C) together to receive a reward, making the task more challenging and likely affecting their performance. Overall, the findings indicate that i.n. administration of 5 mg/kg with 25% w/w CBD-loaded F127 Pluronic^®^ F127 polymeric micelles over 18 days enhances SC in ASD-like rats and are associated with the significant increase in the CBD bioavailability in the brain after this administration strategy when compared to the oral one (**Table 1**). Despite receiving a higher dose, the OR+ group demonstrated a comparatively lower increase in cooperation, suggesting that oral administration is less efficient in delivering CBD to the brain. Meanwhile, i.n.- and the untreated Low rat groups showed no significant changes, confirming that the improvement observed can be attributed to CBD treatment.

**Figure 4.**
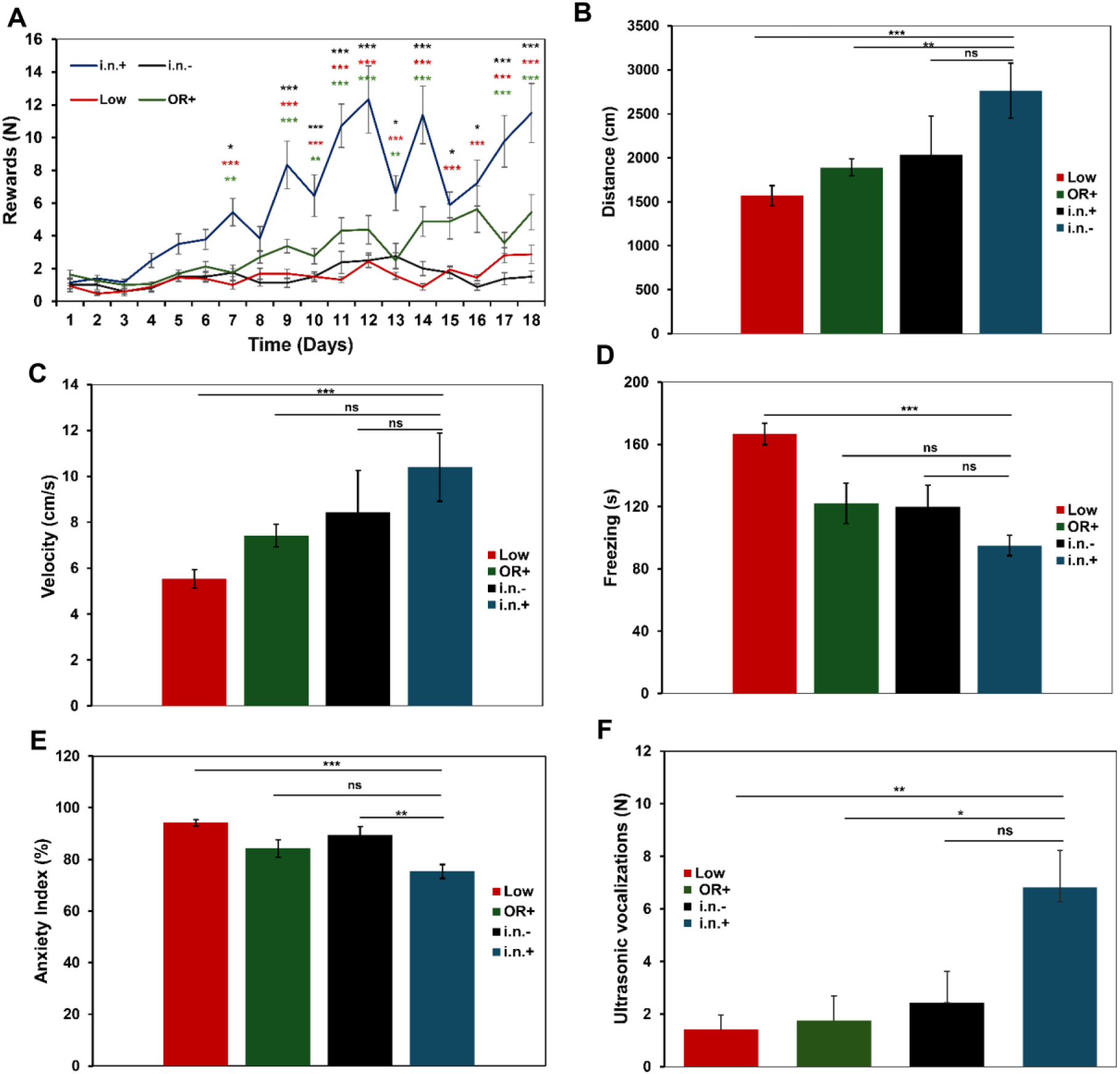
Behavioral tests following treatment of the ASD-like rats. (A) SC test in rats for four treatment groups, i.n. administration of 5 mg/kg of CBD-loaded Pluronic^®^ F127 polymeric micelles (n = 36), oral administration of 15 mg/kg of CBD-loaded Pluronic^®^ F127 polymeric micelles (n = 36), 5 mg/kg of i.n. administration of empty Pluronic^®^ polymeric micelles (n = 18) and untreated group (n = 32). (B-E) Open field test of i.n.+ administration of 5 mg/kg of CBD-loaded Pluronic^®^ F127 polymeric micelles (n = 8), oral administration of 15 mg/kg of CBD-loaded Pluronic^®^ F127 polymeric micelles (n = 8), 5 mg/kg of i.n. administration of empty Pluronic^®^ polymeric micelles (n = 8) and untreated group (n = 24). (F) Number of ultrasonic vocalization of rats after four treatment group i.n. administration of 5 mg/kg of CBD-loaded Pluronic^®^ F127 polymeric micelles (n = 16), oral administration of 15 mg/kg of CBD-loaded Pluronic^®^ F127 polymeric micelles (n = 18), 5 mg/kg of i.n. administration of empty Pluronic^®^ polymeric micelles (n = 14) and untreated group (n = 22). All data are presented as mean ± S.E. ns: Non-statistically significant differences. * Statistically significant difference (p < 0.05); ** statistically significant difference (p < 0.01) and *** statistically significant difference (p < 0.001).

To evaluate the effect of i.n. CBD administration on other core symptoms of ASD, we conducted the open field test to evaluate locomotor activity and anxiety on the same groups described above (n = 8 for i.n+, OR+ and i.n.- groups; 4 female and 4 male, and n = 24 for the untreated rats; 12 female and 12 male). The locomotor activity test is a behavioral test used to assess the spontaneous movement and exploration of animals in a novel environment. It measures the total distance traveled, velocity, and duration of movement, among other parameters. Velocity can be interpreted as a measure of intrinsic motivation and can be affected by various treatments or conditions.^78^ The i.n.+ group showed the highest locomotor activity compared to the other groups, with an average of 2763 ± 311 cm (**Figure 4B**). Conversely, the i.n.- group’s, rats moved a distance of 2032 ± 444 cm, while those of the OR+ moved a distance of 1891 ± 97 cm. The untreated Low rat group showed the lowest activity and moved a distance of 1579 ± 114 cm. These results suggest that the i.n. administration of 25% w/w CBD-loaded Pluronic^®^ F127 polymeric micelles (CBD dose of 5 mg/kg) significantly enhances locomotor activity in the open field test in comparison with the oral (p < 0.01) and the untreated group (p < 0.001). Likewise, the i.n.+ group demonstrated the highest velocity (10.4 ± 1.4 cm/s), while the OR+ demonstrated velocity of 7.4 ± 0.4 and the i.n.- group demonstrated velocity of 8.4 ±1.8 and significant increase in comparison to the untreated group that demonstrated velocity of 5.5 ± 0.4 (p < 0.001). These results suggest that the i.n. treatment enhances locomotor velocity in ASD-like rats (**Figure 4C**).

The ‘freezing’ test is a non-conditioned test aimed at evaluating anxiety-like behavior.^79^ The results for freezing behavior (measured in seconds) in the open field test showed distinct differences among the experimental groups. As expected, the untreated group exhibited the longest freezing duration (∼167 s), indicating a higher anxiety (**Figure 4D**). In contrast, the i.n.+ group demonstrated the lowest freezing duration (∼94 s) highlighting a statistically significant decrease (p < 0.001) compared to the untreated group (**Figure 4D**). The results from the anxiety index in the open field test revealed that the untreated group showed the highest anxiety index (94%) similar to the i.n- group (90%), followed by the OR+ group (84%), and ending up with the i.n.+ group that showed lowest anxiety index (75%), representing a significant decrease compared to the i.n.- (p < 0.05) and the untreated groups (p < 0.001) (**Figure 4E**). These findings suggest that both the treatment and its i.n. delivery may have a pronounced anxiolytic effect compared to the other groups.

The ultrasonic vocalization test reflects an additional ASD-like behavior.^80^ It comprises recording and analyzing the high-frequency sounds, which are beyond the range of human hearing, that rodents emit and is utilized to study communication and the effects of treatments.^81^

The results of the ultrasonic vocalization test showed that the i.n.+ treatment (n = 16, 8 female and 8 male) significantly led to higher number of vocalizations (6.8 ± 1.4) in comparison to the OR+ group (n = 18, 12 female and 6 male) that showed 1.8 ± 0.9 (p < 0.05), and the untreated group (n = 22, 11 female and 11 male) that showed 1.4 ± 0.6 vocalizations (p < 0.01). The i.n.- group (n = 14, 8 female and 6 male) also showed a lower number of vocalizations, 2.4 ± 1.2 (**Figure 4F**). These results suggest the improvement in the communication behavior between the ASD-like rats treated i.n. with 25% w/w CBD-loaded polymeric micelles. Finally, we conducted two additional behavioral tests. The object recognition test assesses the ability to discern between familiar and novel objects.^82^ In this test, rats were divided into the same above-described treatment groups and showed similar performance. The i.n.+ group (n = 8 rats, 4 female and 4 male) showed novel preference (the animal’s tendency to spend more time exploring a novel object compared to familiar ones) of 14.4 ± 13.5%, the OR+ group (n = 8, 4 female and 4 male) showed novel preference of 22.0 ± 9.6%, the i.n.- group (n = 8 rats, 4 female and 4 male) showed novel preference of 21.9 ± 8.7% and the untreated group (n = 24 rats, 12 female and 12 male) showed a novel preference of 19.3 ± 10.5%. Results showed no significant differences between the groups, strongly suggesting that CBD does not improve selective attention in the ASD-like rats (**Supplementary Figure S3A**). Similarly, in the Morris water maze test, i.n+ (n = 6, 3 female and 2 male), OR+ (n = 7, 4 female and 3 male), i.n.- (n = 4 rats, 4 female and 3 male), untreated groups (n = 20, 10 female and 10 male) showed comparable performance in the four days session. Results did not demonstrate a consistent progression (either increasing or decreasing) in latency to reach the platform (in seconds) over the four testing days, suggesting that CBD had no significant effect on spatial learning (**Supplementary Figure S3B**) which is not unexpected because, as in other pharmacological interventions attempted in ASD, the therapeutic efficacy is manifested especially in some of the core behavioral symptoms but less on the cognitive deficit.

## 4. Conclusions

Intranasally administered drugs have gain increasing attention owing its great clinical potential in a variety of CNS diseases and disorders.^83,84^ This study demonstrates that intra-nasal delivery of CBD using Pluronic^®^ F127 polymeric micelles as transmucosal nanocarrier offers a promising strategy for enhancing brain bioavailability and ameliorating the core ASD-like behaviors. The optimized nanoformulation achieved high CBD loading (25% w/w) with suitable size and physicochemical properties for i.n. administration. *In vitro* release studies showed a profile described by the Peppas– Sahlin model, indicating combined diffusion and polymer relaxation mechanisms. The formulation exhibited good cell compatibility and permeability across nasal epithelial cells, supporting its potential for an effective i.n. delivery. *In vivo* imaging and pharmacokinetic studies confirmed rapid and efficient brain uptake of CBD following i.n. administration, with higher brain concentrations and bioavailability compared to a 3-fold greater oral dose. Behavioral tests revealed that lower dose of i.n. CBD-loaded micelles significantly improved SC, reduced anxiety-like behavior, and enhanced communicative ultrasonic vocalizations in ASD-like rats. These results highlight the clinical therapeutic potential of CBD in ASD and pave the way for the application of this alternative minimally invasive administration route to tackle the core symptoms of this neurodevelopmental disorder.

## CRediT authorship contribution statement

**R. Awad:** Data curation, Methodology, Investigation. **S. Aga-Mizrachi:** Methodology, Data curation, Methodology. **I. Maoz:** Data curation. **Lilach Simchi:** Methodology. **A. Avital:** Conceptualization, Methodology, Validation, Data curation, Funding acquisition, Project administration, Supervision, Validation, Writing – original draft. **A. Sosnik:** Conceptualization, Methodology, Validation, Data curation, Funding acquisition, Project administration, Supervision, Validation, Writing – original draft.

## Declaration of competing interest

The authors declare that they have no known competing financial interests or personal relationships that could have appeared to influence the work reported in this paper.

### Ethics approval

All experiments involving animals were conducted in accordance with the ethical standards and guidelines set by the Institutional Animal Research Ethical Committee at the Technion – Israel Institute of Technology and the approved protocol #IL080-05-23. All authors complied with all relevant ethical regulations.

## Supporting information

Fig. S1 Description of behavioral tests; Fig. S2: Cumulative CBD release in vitro; Fig. S3: Behavioral tests in ASD rats

NTA video of CBD-loaded polymeric micelles

## Acknowledgments

This work was funded by the NEVET Nanotechnology Grant of the Russell Berrie Nanotechnology Institute (RBNI) at Technion – Israel Institute of Technology (Israel). A.S. thanks the support of the Tamara and Harry Handelsman Academic Chair.

