## Supplementary material for "Nose-to-Brain Administration of Cannabidiol-Loaded Polymeric Micelles Improves the Core Behavioral Symptoms of Autism Spectrum Disorder": Fig. S1 Description of behavioral tests; Fig. S2: Cumulative CBD release in vitro; Fig. S3: Behavioral tests in ASD rats

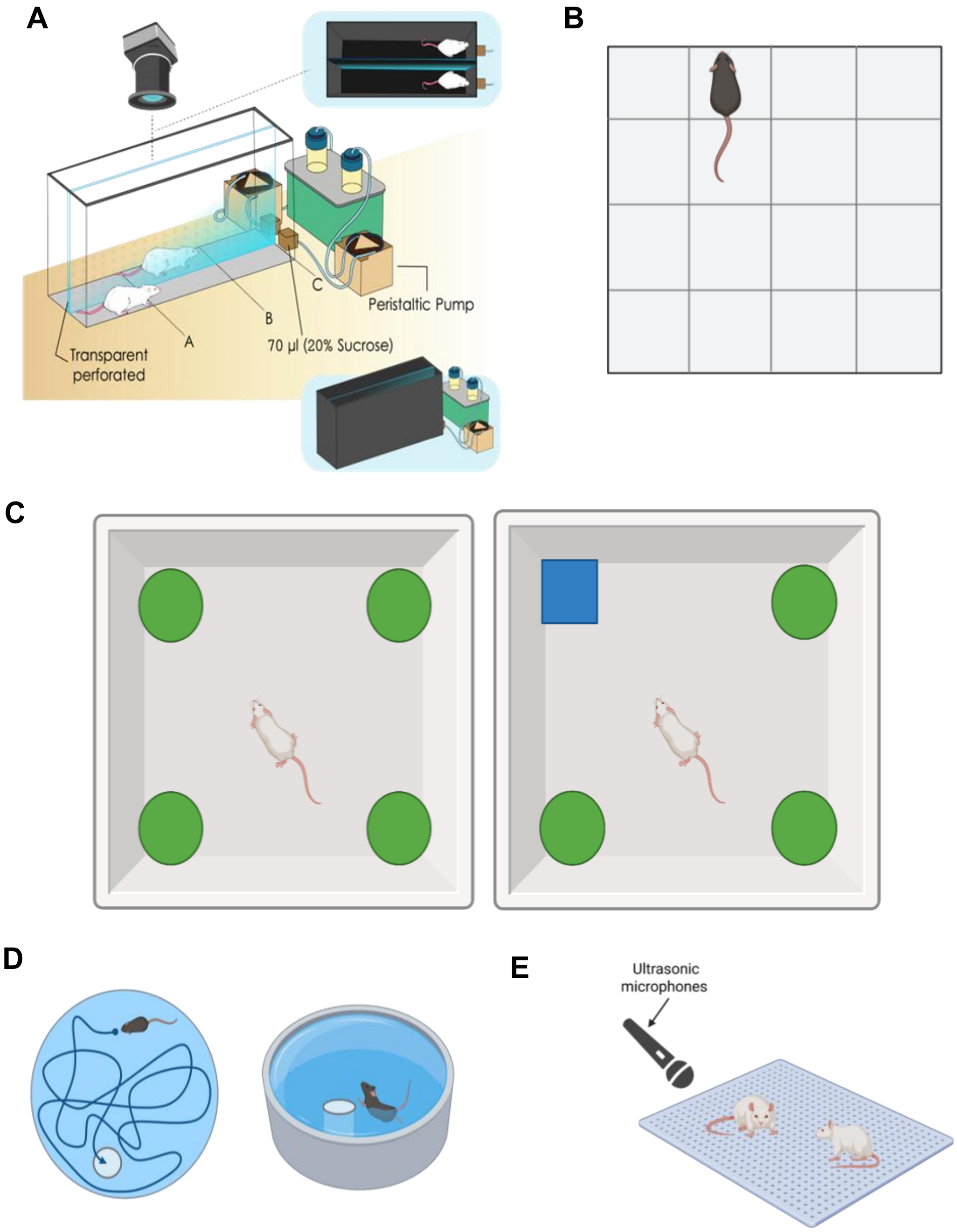


**Figure S1.** Scheme of the different behavioral tests. (A) Social cooperation maze, (B) open field test, (C) object recognition test, (D) water maze test and (E) ultrasonic vocalization test.

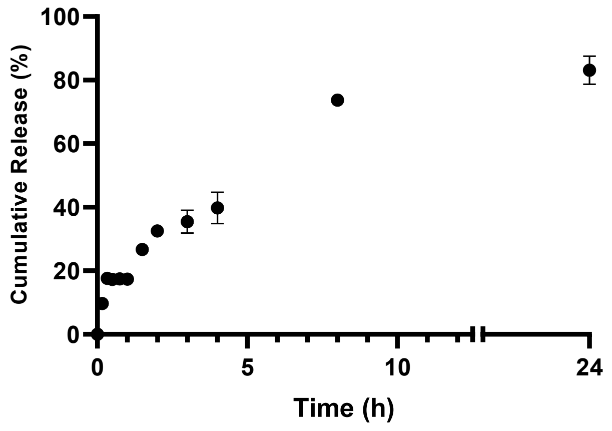


**Figure S2.** Cumulative release of CBD from 25% w/w CBD-loaded Pluronic® F127 polymeric micelles, at 37°C (n = 3). Results are expressed as Mean ± S.D.

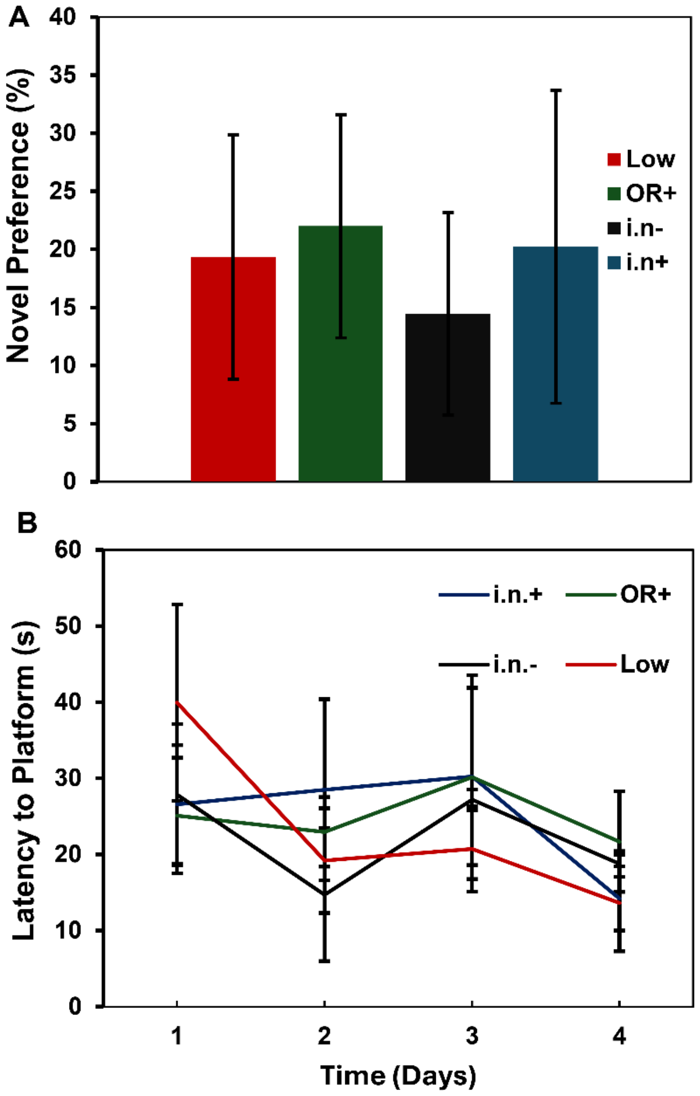


**Figure S3.** Behavioral tests in ASD rats. (A) Novel preference (%) in object recognition test after four treatments group of i.n.+ administration of 25% w/w CBD-loaded Pluronic® F127 polymeric micelles (CBD dose of 5 mg/kg, n = 8), oral administration of CBD-loaded Pluronic® F127 polymeric micelles (CBD dose of 15 mg/kg, n = 8), i.n. administration of empty 10% w/v Pluronic® polymeric micelles (volume of 0.205 μL/g, n = 8) and untreated group (n = 24). (B) Latency to platform in seconds (s) in the Morris water maze test after four treatments group i.n. administration of CBD-loaded Pluronic® F127 polymeric micelles (CBD dose of 5 mg/kg, n = 6), oral administration of CBD-loaded Pluronic® F127 polymeric micelles (CBD dose of 15 mg/kg, n = 7), i.n. administration of empty 10% w/v Pluronic® polymeric micelles (volume of 0.205 μL/g, n = 4) and untreated group (n = 20). Results are expressed as Mean ± S.D.
